## Supplemental figures for "Implicit motor adaptation patterns in a redundant motor task manipulating a stick with both hands"

**Author Contributions:** Conceptualization, Methodology, Investigation, Formal Analysis, Writing: TK, DN, Supervision: DN.

**Keywords:** Reaching movement; Redundancy; Bimanual movement; Motor learning

**Supplementary Figure**

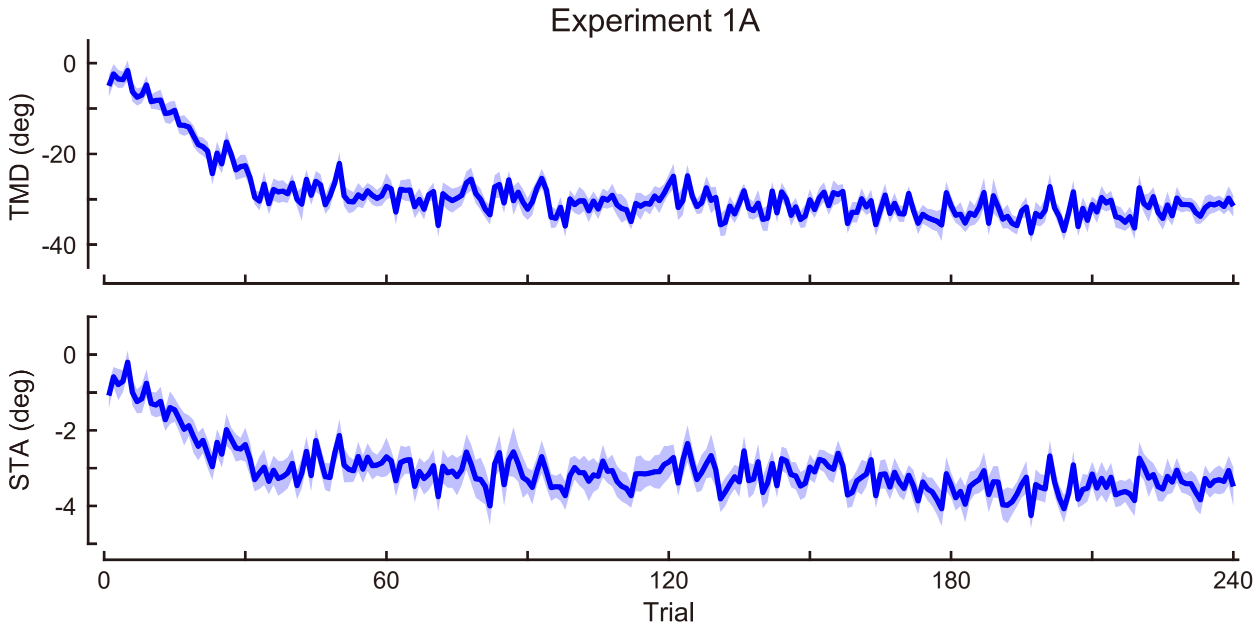


**Supplemental Figure 1. Time-course of the movement pattern in the adaptation phase.** Changes in the movement pattern in the physical space are shown (the tip-movement direction (*top*) and stick-tilt angle (*bottom*)). The data shows mean ± SEM across participants.

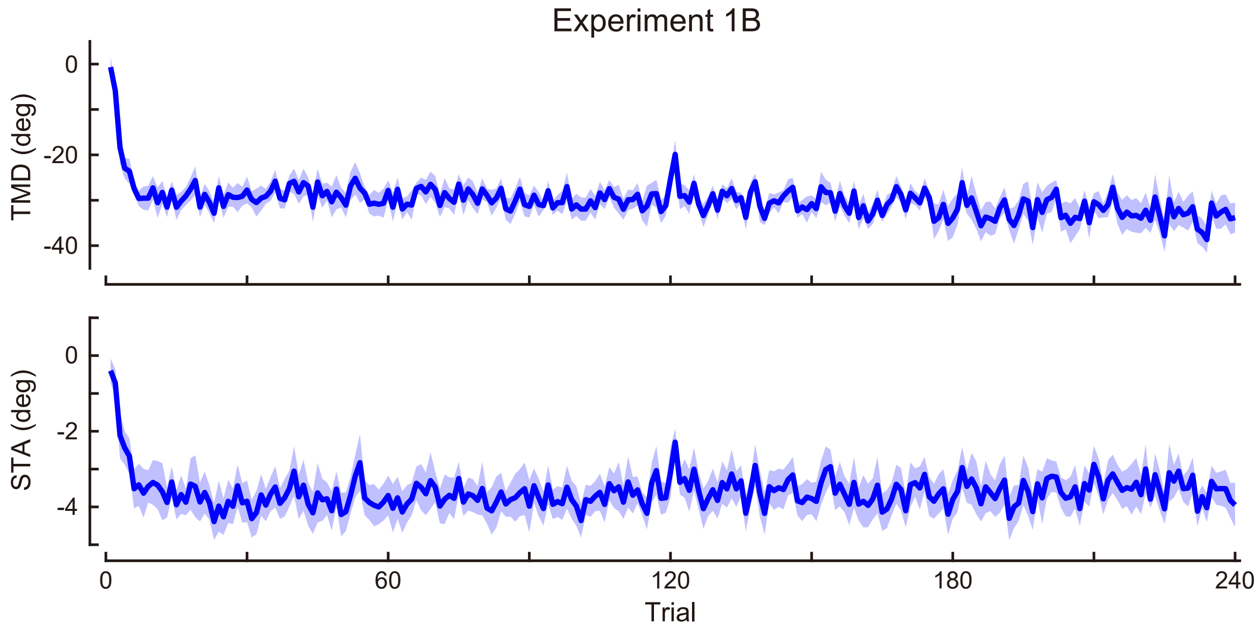


**Supplemental Figure 2. Time-course of the movement pattern in the adaptation phase.** Changes in the movement pattern in the physical space are shown (the tip-movement direction (*top*) and stick-tilt angle (*bottom*)). The data shows mean ± SEM across participants.

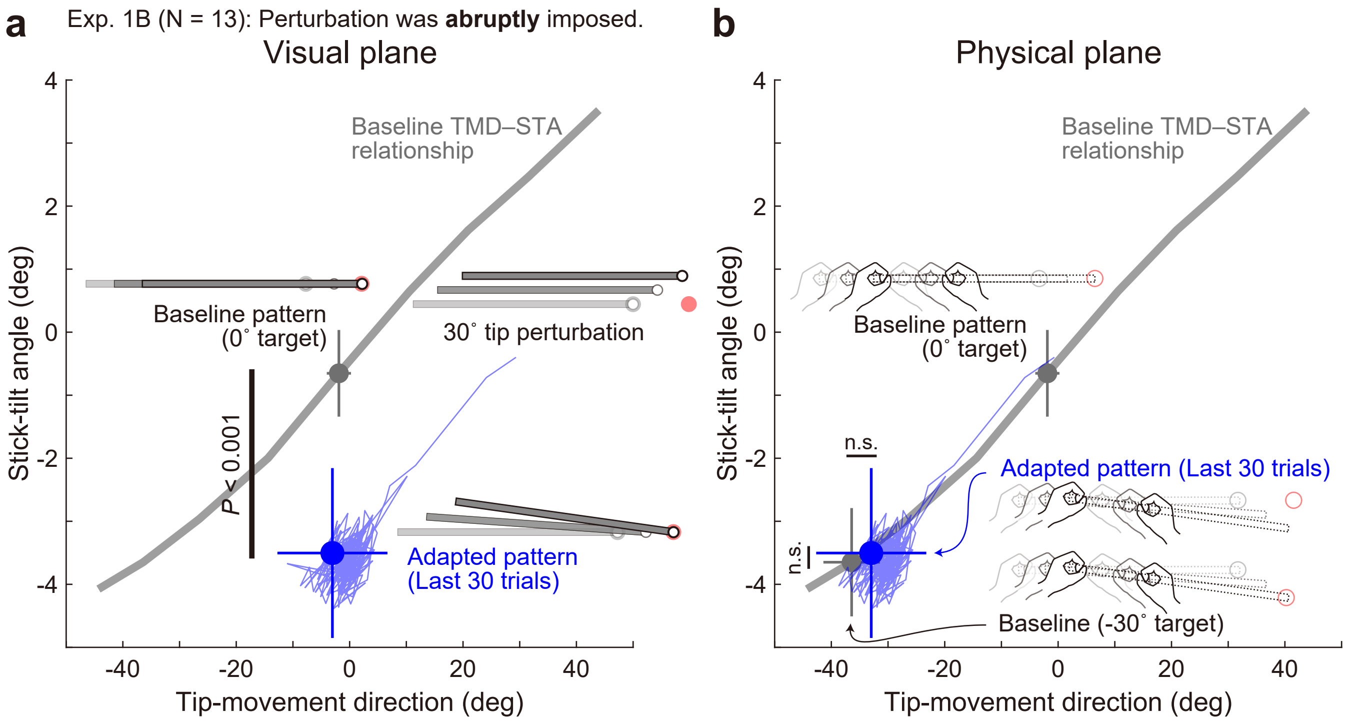


**Supplemental Figure 3. Adaptation patterns to abruptly imposed end-effector relevant perturbations.** (**a**) In Experiment 1B, a 30° tip perturbation was applied from the first trial of the adaptation phase. On the visual plane, the stick-tilt angle after the adaptation (a blue circle) significantly differed from that in the baseline pattern when aiming at the 0° target (a gray circle) (paired *t*-tests, $t(12) = 7.529$, $P < 0.001$), whereas the tip-movement direction remained at the baseline level (paired *t*-test, $t(12) = 0.430$, $P = 0.675$). The formats are the same as Fig. 5a. (**b**) On the physical plane, neither tip-movement direction nor stick-tilt angle in the adapted pattern was significantly different from that in the baseline pattern when aiming at the -30° target (paired *t*-test, TMD: $t(12) = -1.628$, $P = 0.129$, STA: $t(12) = -0.490$, $P = 0.633$).

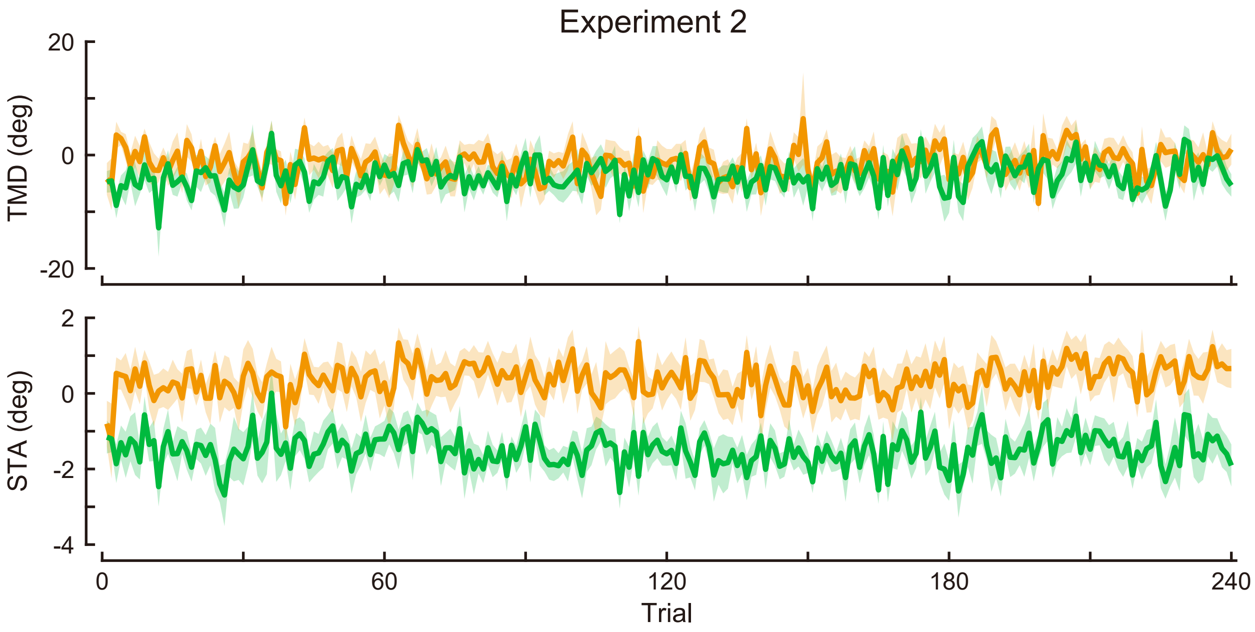


**Supplemental Figure 4. Time-course of the movement pattern in the adaptation phase.** Changes in the movement pattern in the physical space are shown (the tip-movement direction (*top*) and stick-tilt angle (*bottom*)). The data shows mean ± SEM across participants. The green and orange plots indicate E2CCW group and E2CW group, respectively.

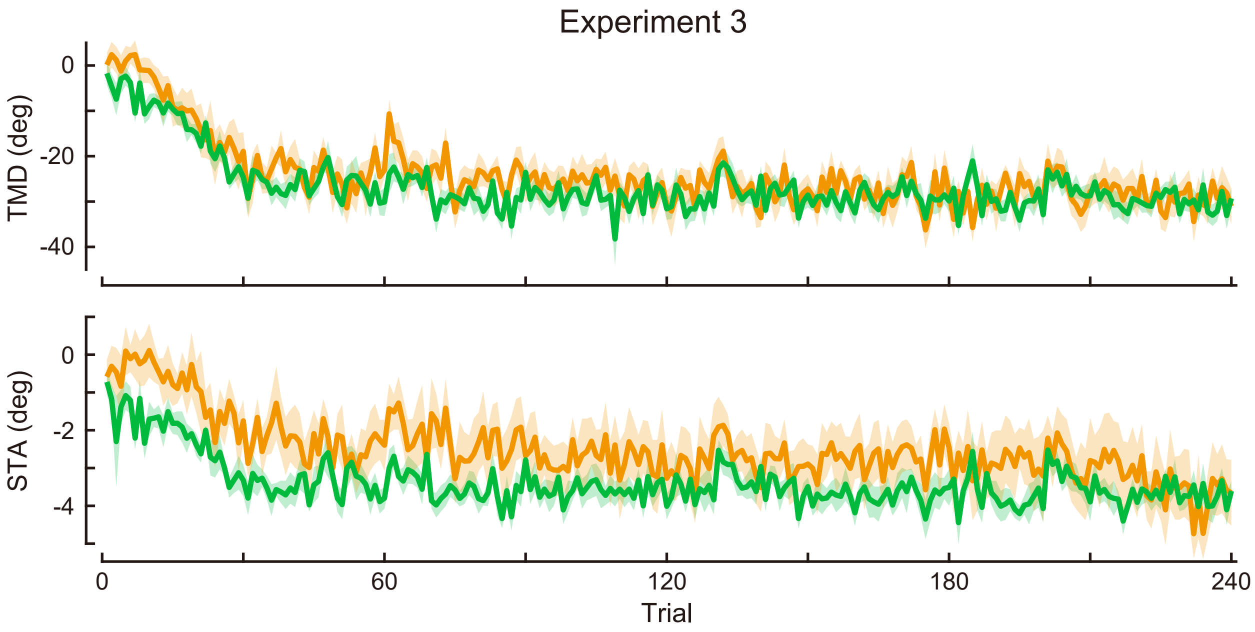


**Supplemental Figure 5. Time-course of the movement pattern in the adaptation phase.** Changes in the movement pattern in the physical space are shown (the tip-movement direction (*top*) and stick-tilt angle (*bottom*)). The data shows mean ± SEM across participants. The green and orange plots indicate E3CCW group and E3CW group, respectively.
